# MADET: A Manually Curated Knowledgebase for Microbiomic Effects on Efficacy and Toxicity of Anticancer Treatments

**DOI:** 10.1101/2022.05.23.493174

**Authors:** Jie Zhang, Xiqian Chen, Jiaxin Zou, Chen Li, Wanying Kang, Yang Guo, Sheng Liu, Wenjing Zhao, Xiangyu Mou, Jiayuan Huang, Jia Ke

**Author notes:** Correspondence should be addressed to J.K., J.H. or X.M. These authors contributed equally to this work (Jie Zhang, Xiqian Chen, Jiaxin Zou).

## Abstract

A plethora of studies have reported the associations between microbiota and multiple diseases, leading to at least four databases to demonstrate microbiota-disease associations, i.e., gutMDisorder, mBodyMap, GMrepo and Amadis. Moreover, gut microbiota also mediates drug efficacy and toxicity, whereas a comprehensive database to elucidate the microbiota-drug associations is lacking. Here we report an open-access knowledgebase, MADET (Microbiomics of Anticancer Drug Efficacy and Toxicity), which harbors 453 manually annotated microbiota-drug associations from 24 papers. MADET provides user-friendly functions allowing users to freely browse, search, and download the data conveniently from the database. Users can customize their search filters in MADET using different types of keywords, including bacterial name (e.g., *Akkermansia muciniphila*), anticancer treatment (e.g., anti-PD-1 therapy) or cancer type (e.g., lung cancer) with different types of experimental evidence of microbiota-drug association and causation. We have also enabled user submission to further enrich the data document in MADET. The MADET database is freely available at https://www.madet.info. We anticipate that MADET will serve as a useful resource for a better understanding of the microbiota-drug associations and facilitate the future development of novel biomarkers and live biotherapeutic products for anticancer therapies.

## 1. Introduction

Gut microbiota plays an important role in carcinogenesis and cancer treatment outcomes [1, 2]. Bacteria including *Helicobacter pylori* [3], *Fusobacterium nucleatum* [4], *Peptostreptococcus anaerobius* [5], enterotoxigenic *Bacteroides fragilis* [6], polyketide synthase-positive (*pks*^+^) *Escherichia coli* [7], and *Campylobacter jejuni* [8], have been reported to contribute to carcinogenesis and tumor development via releasing toxins and activating procarcinogenic signaling pathways [1, 2, 4]. On the other hand, microbes including *Lactobacillus reuteri* [9], *Lactobacillus gallinarum* [10], and *Streptococcus thermophilus* [11] are reported to suppress tumorigenesis and cancer progression via multiple pathways, such as inhibiting the metabolism of tumor proliferation [11]. To date, several databases have been constructed to document these positive or negative associations between gut microbiota and tumor development, such as mBodyMap [12], gutMDisorder [13], GMrepo [14], and Amadis [15]. However, the important effects of microbiota on the efficacy and toxicity of anticancer drugs are not systematically documented.

Gut microbiota modulates the efficacy and toxicity of anticancer drugs through multiple mechanisms including immunomodulation, metabolism, and enzymatic degradation [16]. Notably, the efficacy of immunotherapies (e.g., anti-PD-1 therapy) in cancer patients is positively associated with the abundance of some specific bacterial species (e.g., *Akkermansia muciniphila* [17], *Bacteroides fragilis* and *Bifidobacterium longum* [18]). These commensal gut microbes may reinforce the efficacy of immune checkpoint inhibitors (ICIs) via regulation of the host immune responses (such as increased level of CD4^+^ T cells and/or CD8^+^ T cells in the tumor microenvironment) [17, 19]. Several live biotherapeutic products are under development for their synergistic anticancer effects with immunotherapies, and the clinical trials based on these bacterial products are ongoing, including CBM588 (a *Clostridium butyricum* strain) for treating advanced kidney cancer [20]. Moreover, toxicities of anticancer treatments are impacted by gut microbiota. For example, chemotherapeutic agent irinotecan may be metabolized by gut bacterial β-glucuronidase (which can be found in four major bacterial phyla: Bacteroidetes, Firmicutes, Verrucomicrobia, and Proteobacteria) into its toxic form SN-38, and therefore may inflict toxicity in the gastrointestinal tract, such as epithelial damage and diarrhea [21]. Another example is that immune-related adverse effect (irAE) of ICI therapy, most commonly colitis, has been reported to be associated with the abundance of the Bacteroidetes phylum in the human gut [22]. In a nutshell, microbiota could be employed as a potential biomarker for predicting treatment outcomes and an interventional target for improving the effectiveness of cancer therapies. We therefore argue that with the accumulating evidence of the important roles of microbiota in anticancer treatments, including chemotherapeutic agents, radiotherapy and immunotherapies, it is urgent to develop an open-access and user-friendly knowledgebase to allow for documenting, retrieving and sharing experimental data of these microbiotas and their effects on the efficacy and toxicity of anticancer drugs.

To this end, in this study we implement a knowledgebase, termed MADET (Microbiomics of Anticancer Drug Efficacy and Toxicity; freely available at https://www.madet.info), to provide the most up-to-date manual curation of reports on the interplays between microbiota and anticancer drugs for the researchers in microbiology, pharmacology and other related areas (**Figure 1**). To the best of our knowledge, MADET is the first database providing experimental evidence of associations between microbiota and anticancer drugs with user-friendly functionalities, including data download and flexible keyword search with bacterial name, treatment, or cancer type. We anticipate MADET will serve as a steppingstone for the development of novel microbiomic biomarkers for predicting the outcome of anticancer drugs and live biotherapeutic products for improving the outcome of anticancer drugs.

**Figure 1.**
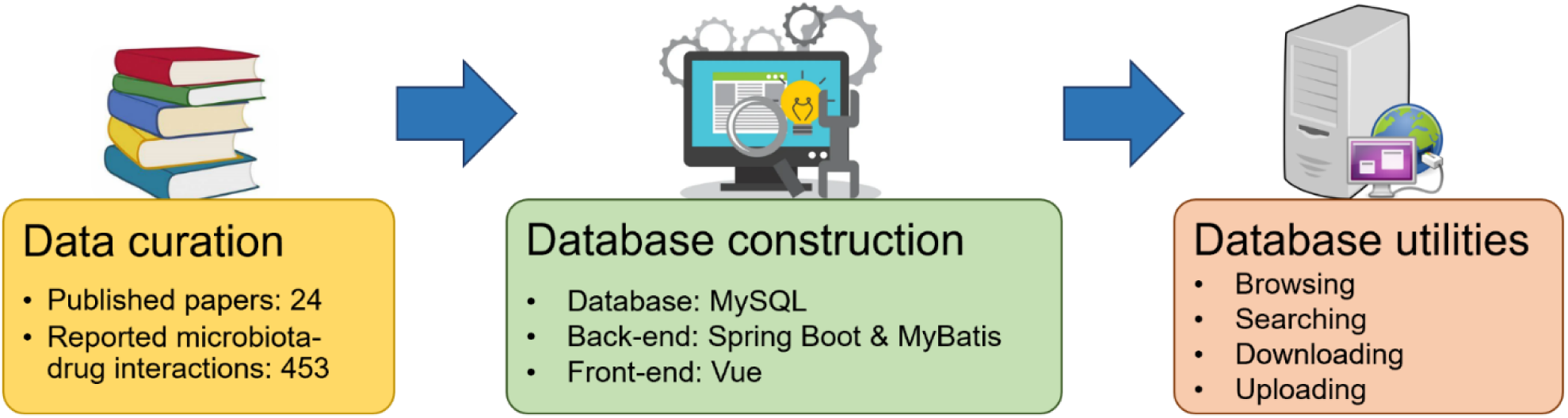
The framework of MADET, including data curation, database construction and utilities.

## 2. Methods

### 2.1. Data collection

To collect the state-of-the-art scientific reports on pharmacomicrobiomics of anticancer drugs, we searched the literature using the keywords “microbiome” and “cancer treatment (e.g., 5-FU, anti-PD-1 or radiotherapy, etc)” on PubMed and extracted 24 published papers involving how the composition of microbiota is associated with the efficacy and toxicity of anticancer drugs. Fourteen of the 24 papers focused on the associations between microbiota and ICIs, and six papers examined the effects on chemotherapeutic agents and their combinations (**Figure 2A**). We further included four publications on cancer radiotherapy studies, given the fact that radiotherapy has been widely applied to cancer treatment. In total, 453 associations between microbiota and drug outcome have been manually identified and recorded in MADET.

**Figure 2.**
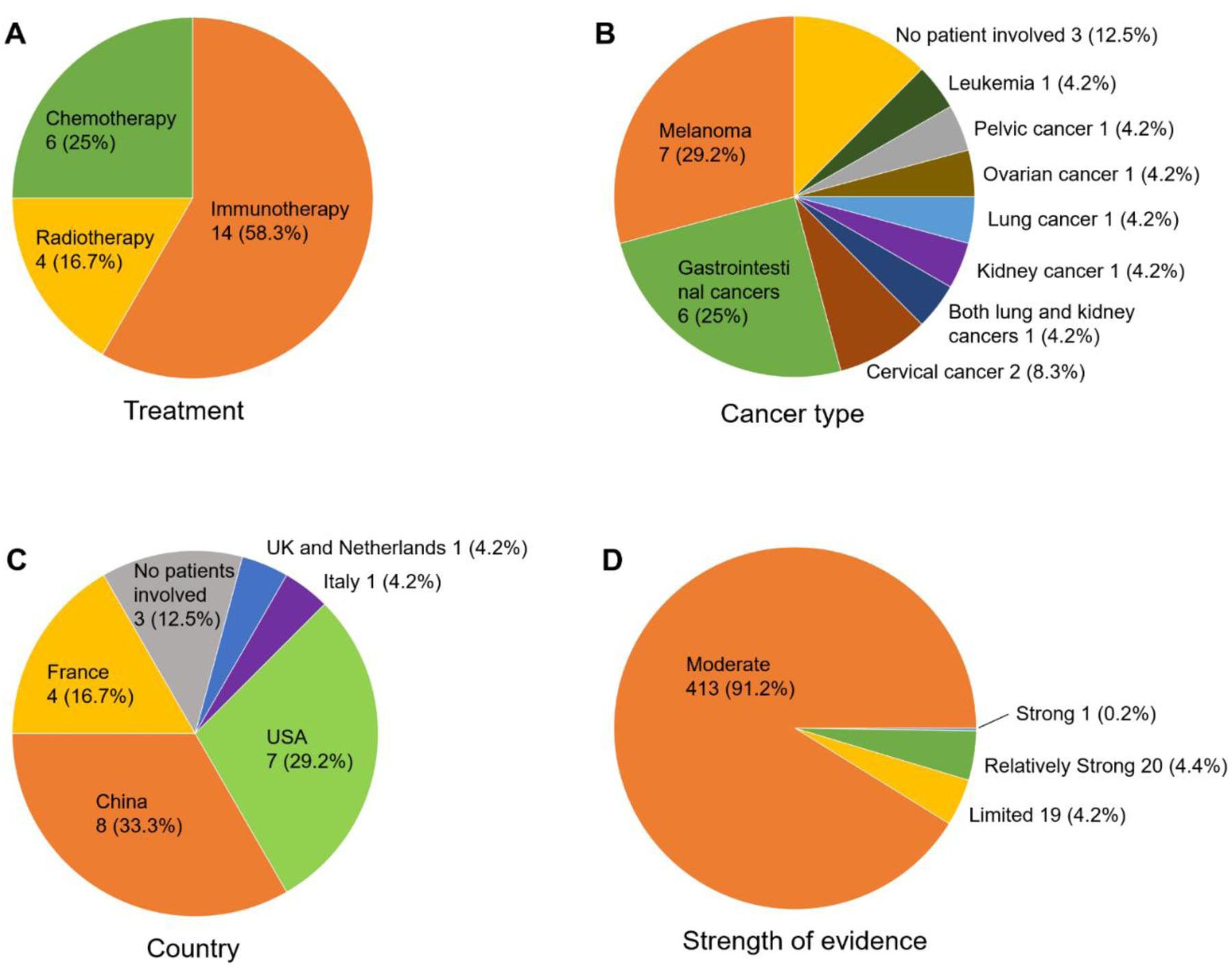
Statistics of curated studies for microbiota-drug associations. The collected studies have been categorized by (A) treatment approaches, (B) cancer types (one study recruited both kidney cancer patients and lung cancer patients), (C) countries where the study was conducted, and (D) the strength of evidence.

We also analyzed our collected studies based on the cancer type and the country where the research was conducted (**Figure 2BC**). Melanoma and gastrointestinal cancers attracted major attention from anti-cancer pharmacomicrobiomics researchers, which comprise 7 and 6 papers, respectively. We further set up several criteria in terms of experimental types for categorizing the collected 453 microbiota-drug associations: “Limited” (only with animal data); “Moderate” (clinical observation of association); “Relatively strong” (clinical observation of association and experimental validation of causation using animal model); and “Strong” (clinical observation of association and validated of causation by clinical trial). Among those, the majority (413 associations) have a “Moderate” strength of evidence (**Figure 2D**). Only one association was labeled as “Strong”: a *Clostridium butyricum* strain CBM 588 increases the effect of immunotherapy (nivolumab and ipilimumab) in treating patients with kidney cancer, which was validated by a Phase 1 clinical trial [20].

### 2.2. MADET construction

MADET was developed and implemented utilizing widely applied open-source packages including Vue (front-end), Spring Boot and MyBaits (back-end), and MySQL (relational database design), to provide reliable delivery of a complex scalable web application (**Figure 1**). The modern and separate front-end/back-end architecture of MADET allows collaboration between developers with different specializations and simplifies the upgrading process for future improvement. The main proxy uses a high-performance Nginx server (https://www.nginx.com) to communicate between the front-end and the back-end programs. The front-end of MADET was constructed with a commercial-used open-source Vue.js framework (https://vuejs.org) and Element UI Toolkit (https://element.eleme.io) for the design of a professional user interface experience design. The back-end of MADET has been designed to be a microservice architecture software, which is facilitated by Java Spring Boot (https://spring.io/projects/spring-boot). The microservice architecture implemented three independent functions modules (services), i.e., the search, browse, and upload services. This highly modular architecture allows MADET to provide software as a service (SaaS) and the possibility of turning MADET into a mobile app and adding new services in the future. The actual data stored in MySQL (sql) can be easily converted to other data formats such as xlsx (Microsoft Excel) and csv (comma separated values). The MADET website resides on Tencent Cloud to leverage the burden of various website configurations and security measures while preserving full control of the web application and data integrity to MADET developers.

## 3. MADET utilities

MADET provides four major functionalities, including database browsing, searching, downloading and user submission with a user-friendly interface (**Figure 3**). The *Homepage* contains two sections. The top section provides a brief introduction to the MADET database and the generic statistical information, such as treatment and cancer type, of the current MADET version using interactive pie charts. Users can click on each pie chart to be redirected to corresponding result pages. For example, if “China” from the pie chart is clicked, MADET will show filtered search results that contain studies conducted in China. The Browse page enables users to view all data available on MADET and allows users to set customized filters to shortlist the results. Each MADET entry contains twelve columns, including Bacterium, Treatment, Potential effects on efficacy, Potential effects on toxicity, Cancer type and Reference, Treatment, Potential effects on efficacy, Potential effects on toxicity, and Cancer type. All selected data will be displayed as a table. Several column headers are provided with an arrow, clicking on which will show a drop-down menu of subcategories will appear for users to narrow down the results. The filter will auto-apply when a new attribute is selected, and the table will auto-refresh to display the filtered results. Multiple subcategories from different attributes can be selected at the time for customized filters. If the number of entries exceeds the limit on one page, users can click the pages button at the bottom of the table to scroll through the data. Searches on microbiota-drug interactions of interest is straightforward. Three types of searching are available on the “Search” page, including “Bacterium”, “Treatment”, and “Cancer type”. After obtaining the desired keywords provided by the user, the search engines will only search the database entries with the provided keywords. We have also provided default keyword for each search type. Users can simply click the “Search” button to see the results extracted using the default keywords. The search results are displayed as a table similar to the table on the Browse page. A drop-down menu is provided from “Treatment” and “Cancer type”, respectively, to narrow down the search results of interest. MADET allows users to download its data for high-throughput or computational analysis. Users can download the sql (MySQL) or xlsx format of the data collected by MADET through the download page. To further enrich the data documented in MADET, we have enabled users to submit their own research data via our submission function on the “Upload” page. We request the users to describe the information that should be covered in our database in detail, including treatment, bacterium and cancer type. The submitted data will be manually reviewed by the MADET team prior to data publishing in the database. The users will be contacted if more clarification is needed via their name and email address provided during data submission. We will confirm with the users once their data has been accepted for publishing in the MADET database.

**Figure 3.**
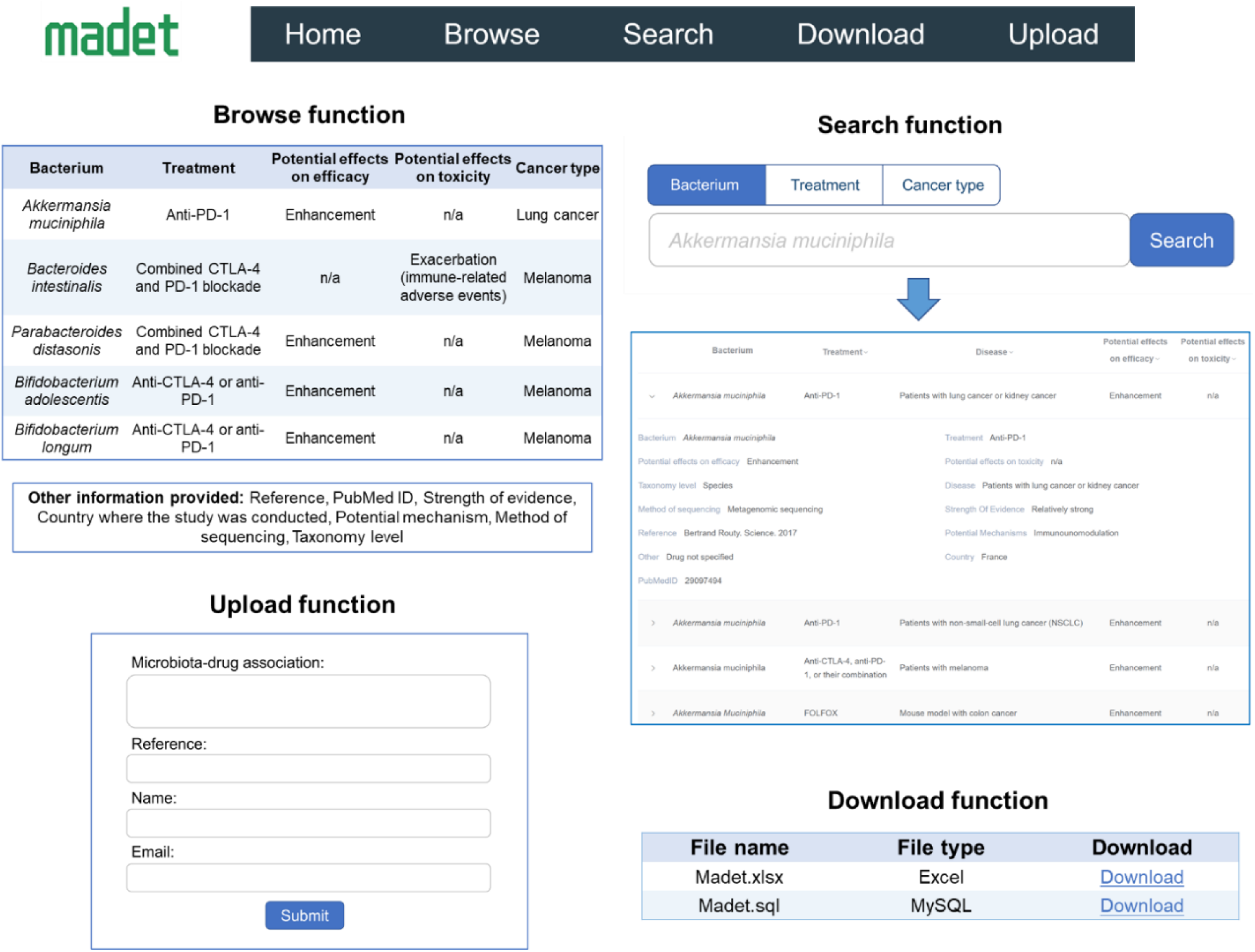
Major functions provided in MADET, including browsing, searching, downloading, and uploading.

## 4. Case study

Some interesting findings involving microbiota-drug associations can be discovered using our MADET database. First, certain bacterial species may exert opposite effects on modulating the efficacy of different drugs. For instance, by searching MADET with the keyword “*Fusobacterium nucleatum*”, we are able to see that the bacterial species *F. nucleatum* is negatively associated with the decreased anticancer effects of 5-FU chemotherapy for colorectal cancer [23] as well as positively related to the promoted efficacy of PD-L1 blockade on the other hand [19]. This suggests that a comprehensive evaluation may be needed prior to the administration of next-generation probiotics targeting microbiota-drug association.

Second, even for the same anticancer drug, we found that same bacterial species showed opposite potential effects against different cancer type or across different countries. For example, searching “*Roseburia intestinalis*” in MADET showed this species is positively associated with the efficacy of immune checkpoint inhibitor (ICI) therapy in patients with lung cancer in France [24], whereas *R. intestinalis* was reported to be negatively associated with the efficacy of ICI therapies in melanoma patients in the USA [18]. Similar results on *Parabacteroides distasonis, Lactobacillus vaginalis*, and *Eubacterium hallii* can be unveiled through searches in MADET with relevant keywords. These findings highlight that the factors including cancer types, geographic locations and analytical methods may affect the conclusion of microbiomic effect on anticancer drugs. Moreover, the functional differences may be strain dependent, i.e., heterogeneous bacterial strains within the same species may have opposite functions on drug outcome. Thus, MADET provides a valuable platform to summarize and analyze the precise mechanism of the observed microbiota-drug associations demonstrated in the current literature.

## 5. Conclusion

With the rapid expansion of our knowledge of gut microbiota during the past decade, multiple bacterial species have been identified as promising candidates for biomarkers of drug susceptibility and live biotherapeutic products for improving drug outcomes. By manually curating 453 associations between microbiota and anticancer drugs with a variety of factors including bacterial species, treatment, cancer type and strength of evidence, the MADET database provides the first comprehensive knowledgebase of the associations between microbiota and anticancer therapeutic outcomes. Users utilize MADET to examine the up-to-date microbiota-drug associations of their bacterium/drug of interest, to advance the generation of novel research hypotheses. MADET will keep evolving with the new data provided by the novel studies on microbiota-drug associations. We anticipate that MADET will serve as a convenient platform for facilitating pharmacomicrobiomics research, including the development of novel biomarkers for predicting drug outcomes as well as novel live biotherapeutic products for improving the outcomes of anticancer drugs.

## Data availability

The MADET database is freely available at https://www.madet.info. Any code or other data related this work will be available upon reasonable request.

## Funding

This work is supported by National Key Research and Development Program of China (2020YFA0907803), Shenzhen Science and Technology Innovation Program (KQTD20200820145822023), and National Natural Science Foundation of China (31900056 and 32000096).

## Disclosure of interest

The authors report no conflict of interest.

## Notes

https://www.madet.info/

### Competing Interest Statement

The authors have declared no competing interest.

